# The structure of a NEMO construct engineered for screening reveals novel determinants of inhibition

**DOI:** 10.1101/2024.07.18.604176

**Authors:** Amy E. Kennedy, Adam H. Barczewski, Christina R. Arnoldy, J. Pepper Pennington, Kelly A. Tiernan, Maria Beatriz Hidalgo, Caroline C. Reilly, Michael J. Ragusa, Gevorg Grigoryan, Dale F. Mierke, Maria M. Pellegrini

## Abstract

NEMO is an essential component in the activation of the canonical NF-κB pathway and exerts its function by recruiting the IκB kinases (IKK) to the IKK complex. Inhibition of the NEMO/IKKs interaction is an attractive therapeutic paradigm for diseases related to NF-κB mis-regulation, but a difficult endeavor because of the extensive protein-protein interface. Here we report the design and characterization of novel engineered constructs of the IKK-binding domain of NEMO, programmed to render this difficult protein domain amenable to NMR and X-ray characterization, while preserving the biological function. ZipNEMO binds IKKβ with nanomolar affinity, is amenable to heteronuclear NMR techniques and structure determination by X-ray crystallography. We show that NMR spectra of zipNEMO allow to detect inhibitor binding in solution and resonance assignment. The X-ray structure of zipNEMO highlights a novel ligand binding motif and the adaptability of the binding pocket and inspired the design of new peptide inhibitors.

## Introduction

The NF-κB Essential Modulator (NEMO) is an essential scaffolding protein in the NF-κB canonical pathway. Sitting downstream of multiple stimuli that initiate the signaling cascade, NEMO assembles key factors in a multiprotein complex that leads to phosphorylation of the kinases IKKα and IKKβ, followed by phosphorylation of the Inhibitor of κB protein (IκB). Subsequent ubiquitination and proteasomal degradation of IκB release the NF-κB dimers to enter the nucleus and activate gene expression. While NF-κB plays a key physiological role in the regulation of many cellular processes, including cell proliferation and survival, B-cell and T-cell maturation, and inflammatory response^1^, many human diseases are driven or sustained by dysregulation of the pathway^2^, counting autoimmune^3^, inflammatory^4^, neurodegenerative^5^, age-related^6^, metabolic^7^ and vascular^8^ diseases and the majority of cancers^9^.

NEMO’s scaffolding function is effected through multiple PPIs^10–12^ engaging domains form the N- to C-terminus, and here we describe our research focusing on modulating the PPI of NEMO with the kinase IKKβ. This interaction is unique to the canonical NF-κB pathway and essential for its activation and therefore represents a prime therapeutic target to develop NF-κB modulators. Specific inhibitors of the NEMO/IKKβ interaction, targeting the IKK-binding domain of NEMO, may avert the poor tolerance observed with inhibition of more pleiotropic players, like kinases or proteasomal inhibitors^13^. Although no crystal structure of full length NEMO has been reported yet, the interaction between NEMO and IKKβ, has been characterized both biochemically^14^ and structurally by X-ray crystallography^15,16^, revealing interaction hot spots that can be exploited by a small molecule or peptide inhibitor for novel therapies.

The C-terminal domain of IKKβ (701-745) binds NEMO with high affinity (15-80 nM)^15,17^ engaging NEMO’s N-terminal sequence 44-111^15^, known as the IKKβ-binding domain or IBD. The structure of the complex of NEMO and IKKβ is a four-helix bundle, where the NEMO coiled-coil dimer opens into a cleft accommodating the two IKKβ helices. The majority of the binding energy is provided by three hot spots on IKKβ identified by Alanine-scanning mutagenesis^14,18^ and consisting of patches of hydrophobic residues: L708/V709 (hot spot 1), V719/I723 (hot-spot 2) and F734-W741 (hot spot 3). IKKβ residues in hot spot 3 correspond to the NEMO Binding Domain (NBD) peptide (fTALDWSW) first described in 2000^19^ for its ability to block the interaction of both IKKs with NEMO *in vitro* and in cells and to inhibit NF-κB activation in cells and *in vivo*. The effectiveness and safety of the NBD peptide as an NF-κB inhibitor has been reported in over 100 publications, for cellular and *in vivo* model systems of different diseases. However, despite the promise of NBD peptides as inhibitors of NF-κB activation, its clinical development was abandoned due to poor bioavailability and low plasma stability^20^. Nevertheless, the inhibition of NEMO PPI provides an exciting opportunity to inhibit NF-κB signaling.

The discovery of small molecule inhibitors of NEMO’s PPIs has been limited by the classical difficulty of disrupting intracellular interactions mediated by tertiary structure and of mimicking the PPI binding epitope^21–23^ and by challenges specific to NEMO, a protein ill-suited to most structural and biophysical methods used to clarify binding mechanism and binding sites and validate inhibitors. Due to the challenges in using structural and biophysical methods to study NEMO, few small molecules inhibitors of the NEMO/IKKβ interaction have been reported and none have progressed to the clinics. Most recent efforts include inhibitors of the NEMO/IKKβ interaction developed through computational methods (structure-based virtual screening^24^ or structure-based pharmacophore^25^) shown to modulate NF-κB activation in cellular assays and *in vivo* models and the natural product Shikonin^26^, which disrupts the NEMO/IKK complex both *in vitro* and *in vivo*. The development of such inhibitors was hindered by the lack of validation of the target or of a mechanism of action, where it was hypothesized that the inhibitor may bind either partner in the PPI or only the formed complex, in absence of information on binding site, binding mode and structure of the complex.

NEMO is a 419-residue long protein, difficult to handle in biochemical, biophysical and structural studies due to its elongated structure, rich in coiled-coil domains separated by flexible regions. The full-length protein was never successfully crystallized and is not suitable for NMR studies. Isolated domains containing the IKK-binding domain of NEMO (44-111 and similar truncations), although characterized in complex with IKKβ(701-745) as described^15^, are disordered and display conformational heterogeneity in the unbound state, which prevents X-ray crystallography or NMR-based studies.

Our and others’ earlier work indicated that the conformationally heterogeneous isolated IKK-binding domain of NEMO can be stabilized in a bioactive conformation by promoting a dimeric coiled-coil structure. Disulfide stabilized constructs^27^ and coiled-coil adaptors stabilized construcs^16,28^ induce a dimeric coiled-coil fold and improve IKKβ-binding activity, *in vitro* and in cells. We recently reported the X-ray structure of the unbound IKK-binding domain of NEMO utilizing the latter construct^16,29^. The coiled-coil adaptors engineered NEMO construct crystallizes easily and enables structure-based drug discovery efforts through NEMO-inhibitor complexes but is not suitable for NMR-based screening assays due to the elongated size of the extended coiled coil, which causes rapid NMR signal relaxation. Although the existing structures of the IBD can facilitate computationally guided drug discovery efforts, they have also highlighted the conformational changes NEMO undergoes upon ligand binding, suggesting an induced fit mechanism, and the presence of multiple putative binding pockets, pointing to the need for better tools for structure-based inhibitor design, screening and validation.

Here we describe a new engineered sequence of the IKK-binding domain of NEMO (zipNEMO), that was obtained through a mutagenesis approach and developed with the goal of aiding the discovery of inhibitors of the NEMO/IKKβ interaction through X-ray crystallography structure determination and protein-detected NMR-based screening, ligand validation and binding site determination. ZipNEMO achieves high affinity IKKβ binding (with a 20-fold improvement over the wild type IKKβ-binding domain) and stabilizes the dimeric and coiled-coil structure. We have determined the high-resolution structure by X-ray crystallography and report the first heteronuclear NMR spectra of the IKK-binding domain of NEMO alone and in complex with IKKβ. Our results open the way for a rational, structure-based drug discovery effort targeting NEMO, where the structure furthers our understanding of the binding determinants and the NMR data enable protein-detected NMR screening, inhibitor validation and binding site determination.

## Results

### Electrostatic and hydrophobic optimization of NEMO improves on nature

We approached the optimization of the IBD region 44-113 of NEMO, responsible for binding IKKβ, with a series of mutations, designed to enhance dimeric coiled-coil propensity and obtain a small protein with improved folding and stability, amenable to screening and biophysical studies, which maintains wild type (WT)-like IKKβ binding. The mutations focused on the protein regions most likely not to impair binding of IKKβ, as identified by Ala scan mutagenesis^14^ and by the NEMO/IKKβ complex structure^15^. Cys54 was also exempt from mutation, as a Cys54 mutation was shown to destabilize the protein and decrease IKKβ binding affinity for NEMO(44-111) and NEMO(1-120)^14^.

The coiled coil is a common structural motif and consists of, in this case, two helices wrapped around each other to form a supercoil^30^. Each helix is characterized by periodic heptad repeats, usually denoted (*a-b-c-d-e-f-g*) in one helix and (*a’-b’-c’-d’-e’-f’-g’*) in the other (Figure 1 A). Residues *a* and *d* are typically non-polar amino acids buried at the interface between the two helices, while *e* and *g* are charged amino acids which contribute to the dimeric coil stability through salt bridges^31^. The IKK-binding domain of NEMO was shown to deviate from this ideal coiled-coil pattern at numerous positions, with a particularly low coiled-coil propensity between residues 66-103^16^. We introduced mutations at coiled-coil positions occupied in NEMO by “non-ideal” residues with two intended stabilization modes: optimization of the knob-into-hole packing by introducing hydrophobic residues in coiled-coil positions *a* and *d,* in place of WT polar residues; optimization of interhelical salt bridges by mutating residues in positions *g* and *e*^32^. The mutations were introduced iteratively in five groups (Figure 1 B,C) clustered by heptad repeat: group 1 for R75V*^d^*, E78R*^g^*(the superscript denotes the position in the heptad); group 2 for T50E*^g^*, L55R*^e^*; group 3 for I71K*^g^*, C76R*^e^*, L80K*^b^*; group 4 for S68L*^g^* and group 5 for R106E*^g^*, E110V*^d^*. At each mutagenesis stage, the constructs were evaluated by circular dichroism (CD), fluorescence anisotropy (FA), and NMR spectroscopy, to monitor respectively structural content and stability, IKKβ binding affinity and quality of the NMR spectra. Mutations that showed improvement in any aspect and preserved or improved binding affinity for IKKβ were kept for the following round of mutations.

**Figure 1.**
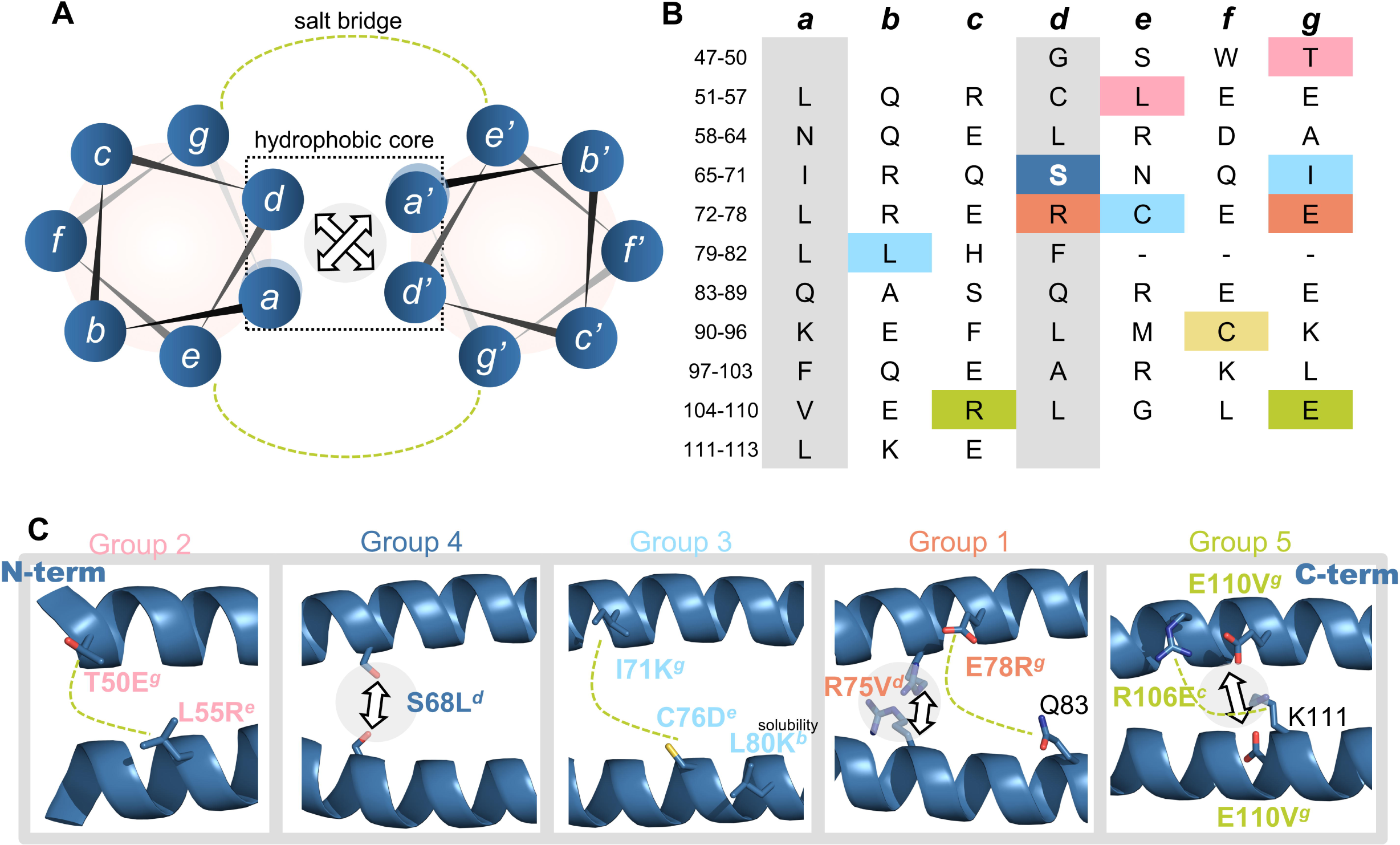
Mutagenesis design. (A) *a*-*g* heptad repeats in a typical coiled coil. Indicated are the *a*/*d* interactions stabilizing the hydrophobic core and *e*/*g* salt bridge interactions. (B) WT-NEMO(44-113) sequence displayed as heptad repeats. Mutated residues are shaded in colors indicating the mutagenesis group. (C) The residues in each mutation group are displayed on the structure of NEMO (3BRV, 6MI3), to highlight the mutagenesis rationale.

CD reports on protein folding, coiled-coil content and protein stability. Far UV circular dichroism spectra of a typical α-helical protein display minima at 222 and 208 nm. Coiled-coil character is identified by a ratio of ellipticity at 222 vs 208 nm greater than 1, while a canonical α-helix displays a ratio smaller than 1^33,34^. The intermediate constructs were examined by CD at five concentrations between 5 and 25 μM, to assess the effect of concentration on coiled-coil folding^35^. The constructs show different coiled-coil propensity but no concentration dependence above 5 μM. Construct stability was assessed by a temperature denaturation experiment monitoring the ellipticity at 208 and/or 222 nm. There was no change in the CD spectrum of each sample acquired before and after the melting experiment, indicating that the unfolding is reversible for these constructs.

The mutation combinations tested included NEMO-1 (the appended numbers indicate the group/s included), NEMO-12, NEMO-124, NEMO-134 and NEMO-125. The best combination of mutations included group 1, 2 and 5 (NEMO-125; see Table 1 for all constructs properties). Of the rejected mutants NEMO-124 (incorporating group 4 S68L*^d^*) aggregated at concentrations above 1 μM and NEMO-134 (incorporating group 3 I71K*^g^*, C76D*^e^*, L80K*^b^*) had decreased affinity for IKKβ. NEMO-125 has improved helical content, coiled-coil content and a 10-fold improvement in affinity for IKKβ over WT-NEMO(44-113) (Table 1 and Figure 2; CD spectra, thermal denaturation curves and binding curves for all mutants are reported in Figure S1). NEMO-125 was further modified to prevent disulfide bond-mediated aggregation at high concentrations, by incorporating the C76S and C95A mutations. Mutation of these two residues was reported to have no effect on NEMO biological activity^36^. NEMO-125CSCA has an improved binding affinity for IKKβ of 272 nM and the best quality NMR spectra with improved line broadening and dispersion, and a number of peaks consistent with a symmetric dimer. This is likely the result of reduced conformational heterogeneity and improved solution behavior due to the lack of higher order oligomers and aggregates. Analysis of ^1^H,^15^N-TROSY spectra of a ^2^H,^15^N labeled NEMO-125CSCA revealed a small number of very intense cross-peaks, which deteriorated spectral quality and could possibly interfere with acquisition of good quality triple resonance spectra for resonance assignment, due to their much slower relaxation rate. We truncated residues N-terminal to the helix-breaking Pro48 to generate NEMO(50-113)-125CSCA and we named it zipNEMO.

**Figure 2.**
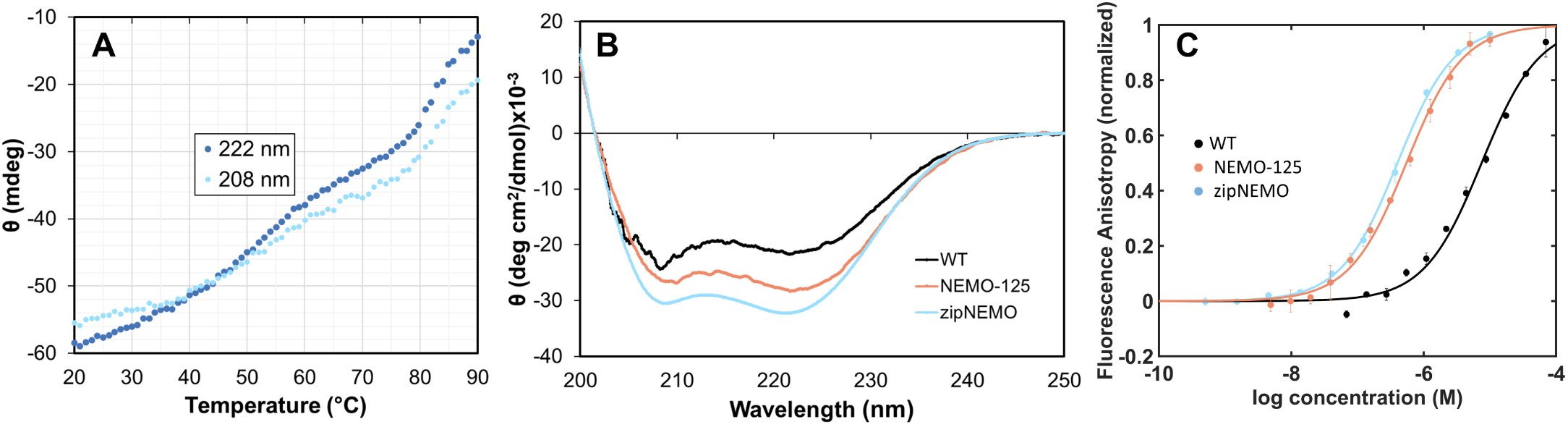
zipNEMO is a stable coiled coil and binds IKKβ with high affinity. (A) Thermal denaturation monitored by CD at 222 and 208 nm. The coiled coil starts unfolding between 39-45°C. (B) Overlay of the CD spectra for the engineered NEMO constructs, showing the stabilization of structure and coiled-coil content compared with WT NEMO(44-113). (C) Binding affinity of the engineered NEMO constructs for FITC-IKKβ_KKRR_(701-745) by Fluorescence Anisotropy, compared with WT NEMO(44-113); lines represent the curve fitting. Error bars represent the standard deviation of 3 repeats. See also Figure S1.

**Table 1.**
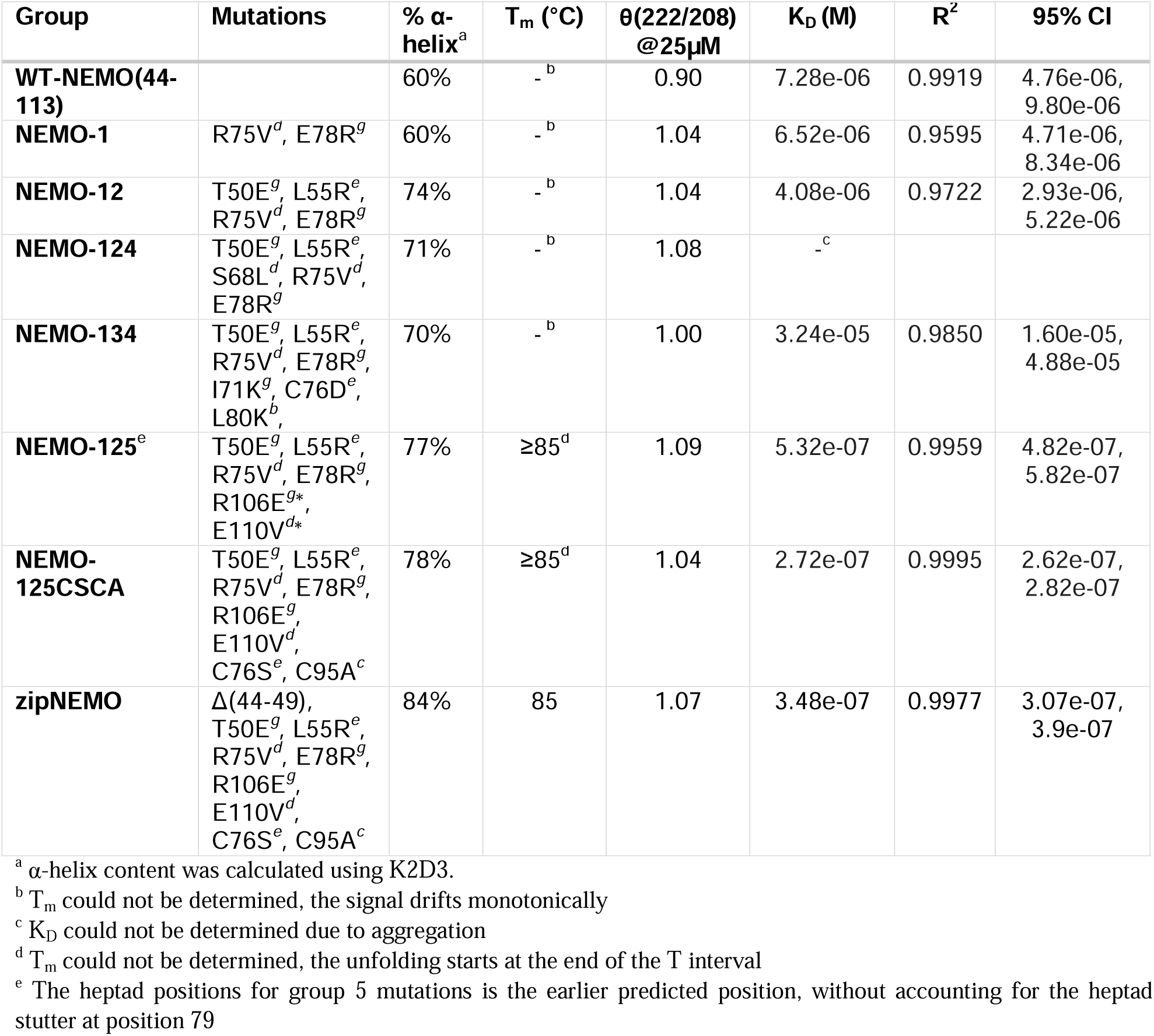
Properties of the engineered NEMO mutants.

### ZipNEMO is a stable coiled coil and binds IKKβ with high affinity

ZipNEMO is 84% helical with a typical coiled-coil ellipticity ratio at 222/208 nm of 1.07 and a cooperative melting behavior with a T_m_= 85°C. The structure maintains coiled-coil character up to 39°C. Above that temperature the structure is still helical but the progressive loss of coiled-coil content indicates the unfolding of the tertiary structure. (Figure 2 A,B). We further investigated the stabilization of the dimeric structure in zipNEMO using reducing agents, cross-linking reagents and SEC-MALS, and compared it to the WT-NEMO(44-113) construct. Both constructs form disulfide mediated dimers (Cys54) which are resistant to reducing agents (TCEP, DTT) as shown by a clear dimer band in SDS-PAGE gels (Figure S2). In presence of a cross-linking reagent (BS2G or BS3) zipNEMO shows the presence of monomer, dimer and tetramer in solutions (Figure S2, BS2G provides the same results as BS3).

The SEC-MALS profile indicates that zipNEMO and WT-NEMO(44-113) elute at similar elution volumes. As it has been reported before^15,28^, NEMO coiled-coil proteins elute at an earlier time than expected for similar MW globular protein, due to the large hydrodynamic radius of the elongated coiled-coil fold. The dispersity of both samples in the MALS experiment is indicated by a trailing shoulder on the peak with decreasing molecular weight, due to the dynamic equilibrium between monomers, dimers and tetramers, with a higher prevalence of dimer/tetramer for zipNEMO (Figure S2). All our experiments indicate the presence of dimeric and tetrameric species in equilibrium in solution. The NMR spectra in phosphate buffer reflect a predominant symmetric dimer in solution, while the crystallization conditions (*vide infra*), favor the tetrameric species.

Stabilization of the dimeric species of the N-terminal domain of NEMO was previously reported to increase binding affinity for IKKβ, for constructs incorporating N- and/or C-terminal ideal coiled-coil adapters^16,28^ and a construct incorporating a L107C mutation^27^. Accordingly zipNEMO binds IKKβ(701-745) with a K_D_ = 348 nM, a close to 20 fold improvement over wild type NEMO(44-113) (K_D_ = 7.3 µM, Table 1 and Figure 2C).

Finally, zipNEMO is the first IBD construct of NEMO to display good quality ^1^H,^15^N heteronuclear correlation spectra that are suitable for resonance assignment, with a number of cross peaks in agreement with a symmetric dimer. The line broadening is consistent with an elongated dimer of this size and indicates a stabilization of the structure and reduction in the conformational exchange observed for the corresponding-length WT-construct (Figure 3A). We monitored binding of unlabeled IKKβ(701-745) to ^15^N-zipNEMO by ^1^H,^15^N-TROSY NMR: Figure 3B shows the extensive chemical shift perturbation originated by IKKβ(701-745) binding to zipNEMO. The result indicates that the construct is suitable for protein-detected NMR-based screening of libraries of ligands. Backbone resonance assignment will allow for the direct identification of ligand binding site by classical Chemical Shift perturbation methods^37^, facilitating the discovery and validation of new specific NEMO /IKKβ inhibitors.

**Figure 3.**
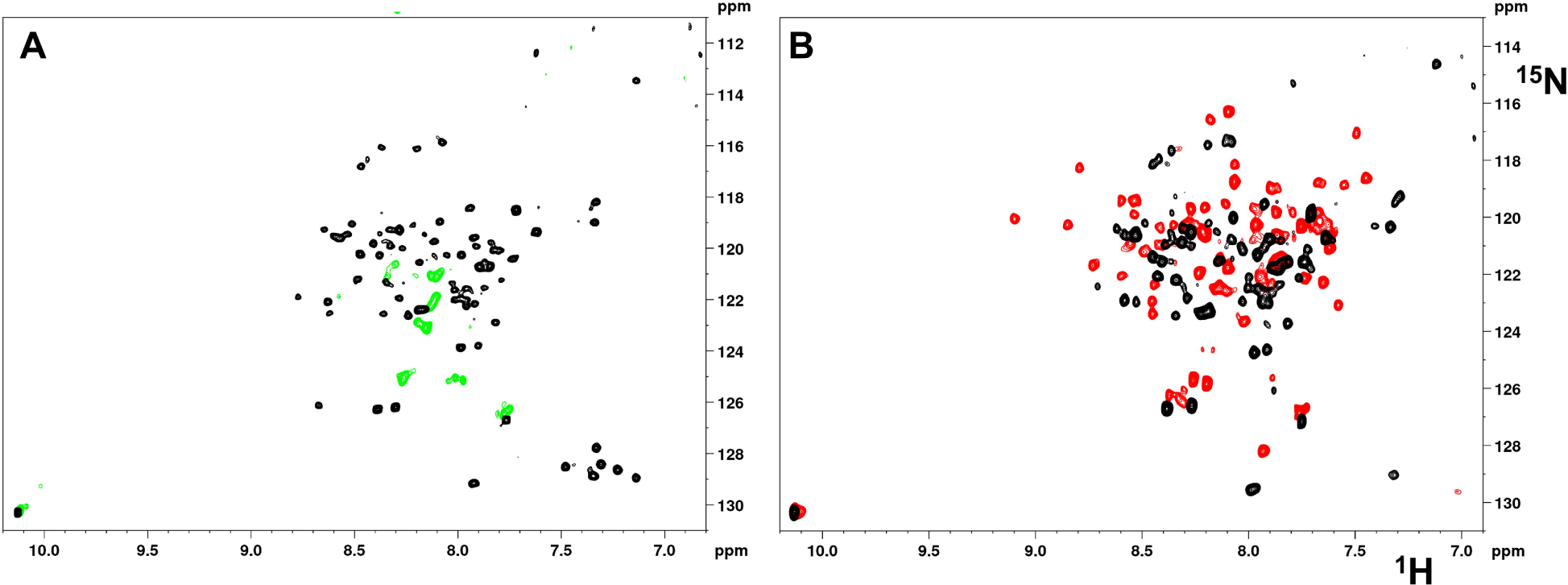
^1^H,^15^N TROSY spectra of ^15^N-zipNEMO. (A) ^15^N-zipNEMO in black compared to ^15^N-WT-NEMO(44-113) in green, which shows only few broadened signals. The zipNEMO spectrum is in HEPES pH 6.5. (B) ^15^N-zipNEMO in black vs. ^15^N-zipNEMO in complex with unlabeled IKKβ(701-745), in red, showing the chemical shift perturbation induced by ligand binding. The spectra are in PBS pH 7.4.

### Longer NEMO constructs incorporating the zipNEMO mutations retain high IKKβ affinity

We further verified the effect of our mutations on a longer construct of NEMO, creating GST-zipNEMO(50-196), which incorporates mutations in Groups 1+2+5, C76S, C95A and the N-terminal truncation. This was chosen as the closest construct to WT-GST-NEMO(1-196), which is our standard for monitoring binding of FITC-IKKβ(701-745) in the FA assay. GST-zipNEMO(50-196) binds FITC-IKKβ(701-745) with a K_D_ = 9.4 nM (R^2^ = 0.997), comparable to WT-GST-NEMO(1-196) (K_D_ = 18 nM, R^2^ = 0.987) (Figure S3) and comparable to what reported for NEMO(1-419) with a K_D_ = 3.6 nM^14^. The experiment validates that the mutations are fully compatible with WT-like binding of IKKβ to NEMO.

### Structure of zipNEMO recapitulates the NEMO/IKKβ complex structure

ZipNEMO crystallized in the P1 space group and the data diffracted anisotropically to 1.878, 1.395 and 1.671 Å. The data was phased by SAD using the anomalous signal of the Lanthanide ions present in the crystallization conditions (PDB: 8U7C, data summary in Table 2). The structure of zipNEMO is a 113 Å long four-helix bundle, formed by two intercalated NEMO dimers, in a relative antiparallel orientation, each dimer stabilized by an interchain disulfide bond at Cys54 (Figure 4 A). Three of the zipNEMO helices adopt a conformation very similar to each other and the helix of NEMO in the NEMO/IKKβ complex structure (3BRV, chain B), with RMSDs between 1.1-1.2 Å. Each dimer is not perfectly symmetric with RMSDs between chains A/C of 1.17 Å and B/D of 1.37 Å. Each dimer is stabilized by a canonical coiled-coil dimerization interface at the N terminus (residues 48-72) which then opens forming a cleft that accommodates the second dimer (Figure 4 A), with helices on both sides of the cleft. The dimerization interface is realized by typical coiled-coil packing of residues in position *a* and *d* at the hydrophobic interface (Cys54, Asn58, Leu61, Ile65, Leu72). The dimer packing validates the mutagenesis design, confirming inter-chain hydrogen bonds involving the side chains of Glu50 and Arg55 (Group-2 mutations: T50E, L55R), stabilizing each dimer at the N-terminus. Additionally, each dimer is stabilized by inter-helix H-bonds involving Asn58^B,D^, Asn58^B^-Cys54^D^_HN_, Glu57^B^-Arg62^D^, Glu57^B^-Asn58^D^ (and corresponding symmetric pairs and A/C pairs; capital letter superscript indicates the chain). All these interactions, with exception of the engineered E50-R55, recapitulate the interactions observed for the WT-NEMO dimer in the NEMO/IKKβ structure (3BRV). Group-5 mutation Glu106 introduces an intra-chain helix-stabilizing H-bond with the side chain of Lys102. The overall architecture of the four-helix bundle combines canonical parallel coiled-coil dimer packing at the N-terminus (50-72) and canonical antiparallel tetrameric coiled-coil packing^38^ in the central region (65-107), where *a* and *d* residues are nearly completely buried at the hydrophobic interface. The structure is also strikingly similar to the four-helix bundle of the NEMO/IKKβ complex, formed by the NEMO dimer and the two bound IKKβ chains (3BRV), where here zipNEMO acts as both the receptor (chains B/D correspond to the NEMO chains B/D in 3BRV) and the pseudo-ligand (chains A/C correspond to the IKKβ chains A/C in 3BRV, Figure 4 B). ZipNEMO chains A/C (the pseudo-ligand dimer in mint in figure 4 B) lie in the cleft formed by chains B/D (the receptor dimer in blue). The NEMO receptor-dimer (chains B/D) adopts a structure almost identical to the WT NEMO dimer in the IKKβ complex (3BRV, chains B/D), with an RMSD of 1.35 Å, with helices overlapping up to residue 97. After Phe97 the zipNEMO helices start to diverge, with interhelical distances increasing to a maximum of around 14 Å at the C-terminus (calculated as (*da* + *da’*)/2 as in Barczewski et al. ^16^; Phe97, Val104), compared to 8.3 Å for NEMO in 3BRV. The opening accommodates the pseudo-ligand-dimer of zipNEMO, chains A/C, which occupy a very similar position to the ligand IKKβ(701-745) in 3BRV (chains A/ C), but extend further due to the increased length and twist into a compact canonical coiled-coil dimer at their N-terminus as described above. The pseudo-ligand-dimer of zipNEMO adopts an overall more open structure, with a maximum interhelical distance of 18 Å (residues 97, 100, 104), slightly larger than the corresponding interhelical distance between the IKKβ chains A/C in the NEMO/IKKβ complex of 16 Å (residues 705, 708, 712). This opening results in an RMSD between zipNEMO dimers B/D vs A/C of 2.63 Å (over 120 residues). For ease of description we will label zipNEMO receptor amino acids (chains B/D) with a three-letter code, and zipNEMO pseudo-ligand amino acids (chains A/C) with a one-letter code, as well as IKKβ amino acids.

**Figure 4.**
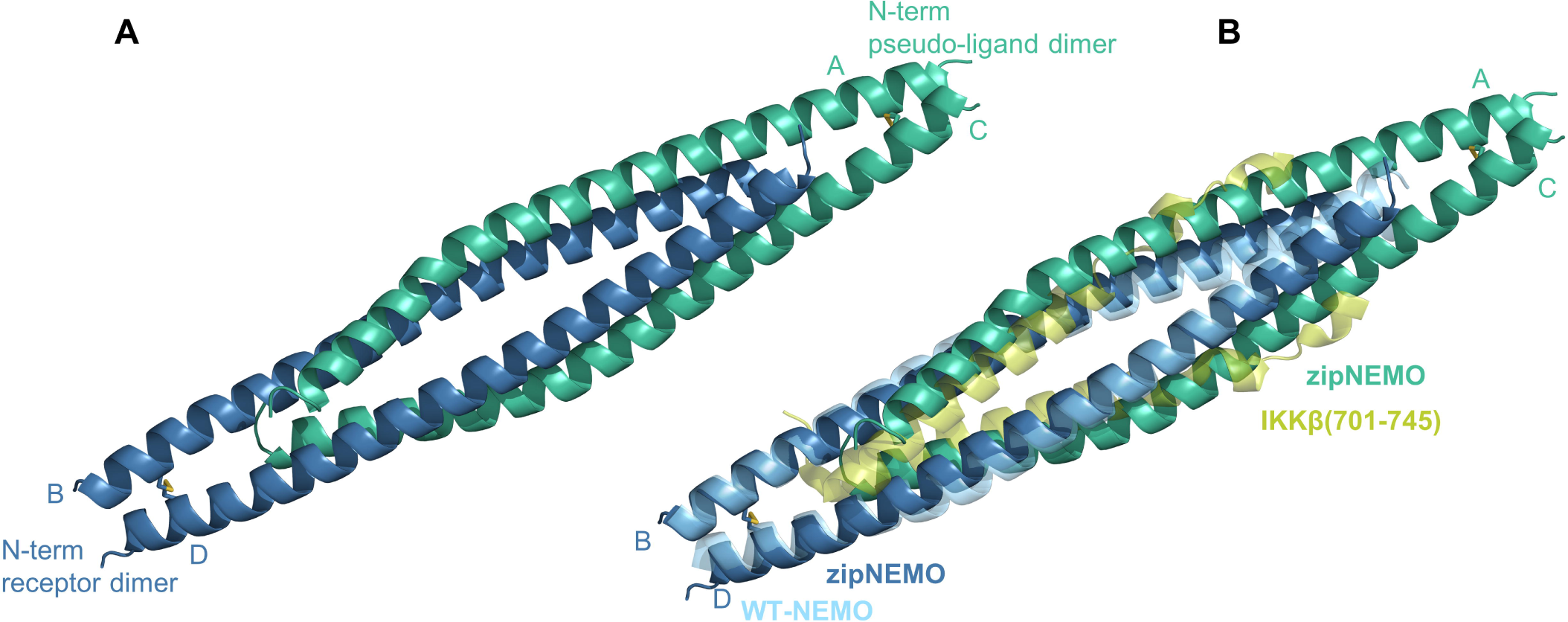
zipNEMO forms a four-helix bundle similar to the NEMO/IKKβ complex. (A) cartoon representation of the structure of zipNEMO: the pseudo-ligand dimer is in mint and the receptor dimer is in blue. The Cys54 disulfide bridges are shown in sticks. The chain names are indicated. (B) superposition of the zipNEMO structure and the NEMO/IKKβ structure (3BRV). The structures were superimposed using the respective chains B/D. IKKβ(701-745) is in pear and NEMO(44-111) is in light cyan (30% cartoon transparency), so that both “ligands” are in shades of green and both receptors are in shades of blue.

**Table 2.** Data collection and refinement statistics for crystallographic experiments.

|  | zipNEMO |  |
| --- | --- | --- |
| **Data Collection** |  |  |
| Space Group | P 1 |  |
| Cell Dimensions |  |  |
| $a, b, c$ (Å) | 37.51, 40.89, 50.15 | |
| $\alpha, \beta, \gamma$ (°) | 92.62, 106.14, 98.87 | |
| Redundancy | 9.4 (6.3)<sup>a</sup> |  |
| Elliptical Completeness % | 91.2 (58)<sup>b</sup> |  |
| $I/\sigma(I)$ | 7.6 (1.7) | |
| $R_{\text{meas}}$ | 0.193 (1.791) | |
| CC $\frac{1}{2}$ | 0.996 (0.426) | |
| Anomalous Completeness (ellipsoidal) | 90.1 (53.1) |  |
| Anomalous Multiplicity | 4.8 (3.3) |  |
| Figure of merit | 0.59 |  |
| Bayesian CC score (x100) | 61.8 +/- 13.7 (2SD) |  |
| **Refinement** |  |  |
| Resolution Range | 47.965 - 1.436<sup>b</sup> (1.614 -1.436) |  |
| Reflections used in refinement | 31669 (9942) |  |
| Reflections used for $R_{\text{free}}$ | 2000 (6.32%) | |
| $R_{\text{work}}/R_{\text{free}}$ | 0.1727/0.2020 | |
| No. of atoms |  |  |
| Protein | 1995 |  |
| Water | 139 |  |
| B factors (Å<sup>2</sup>) |  |  |
| Protein | 23.63 |  |
| Water | 30.08 |  |
| Other | 32.87 |  |
| RMSD |  |  |
| Bond lengths (Å) | 0.759 |  |
| Bond angles (°) | 0.956 |  |

<sup>a</sup> Highest resolution shell is shown in parenthesis
<sup>b</sup> The anisotropic diffraction data was truncated using the STARANISO server to include all valid data (reflections with $I/\sigma(I)$ of 1.2) to resolutions of: 1.878 Å, 1.395 Å and 1.671 Å along the $a^*$ , $b^*$ and $c^*$ axis respectively.

### The hot spots for binding are conserved in zipNEMO

The hot spots for the interaction between IKKβ and NEMO have been identified by Ala-scanning mutagenesis^14^ of IKKβ(701-745) as residues 708/709 (hot spot 1), 719/723 (hot spot 2) and 734-742 (hot spot 3, the NBD peptide sequence). We analyzed the structure of zipNEMO to ascertain whether the three hot spot binding sites are conserved and play a similar role in the formation of the four-helix bundle. In the NEMO/IKKβ complex hot spot 3 contains the strong and highly conserved NBD residues of IKKβ: F734, L737, W739, W741, L742. The pseudo-ligand zipNEMO presents a similar set of hydrophobic residues in the same spatial position. Due to the helical fold, residues L79, V75, L72 and I71 mimic the spacing and orientation of F734, W739, W741 and L742, respectively, in IKKβ (Figure 5 A,B). The hydrophobic cluster, that would be naturally interrupted by NEMO WT residue R75, is here completed by the R75V mutation we introduced in zipNEMO. Similarly to the NEMO/IKKβ complex, we observe two symmetric binding pockets at this site, where the pseudo-ligand dimer chains A/C bind above and below the receptor dimer plane (chains B/D). The hydrophobic pocket is lined on the side by receptor residues Phe92, Leu93, Met94, Phe97 and Ala100, the same NEMO residues engaged by IKKβ^15^, while the opposite pseudo-ligand helix contributes to the bottom plate of the pocket with residues L72, V75, L79, L80. In addition, H-bonds stabilize the interaction: E74^C^/Lys96^D^, R78^A/D^/Glu89^B/C^. The R78 interaction is novel and results from the engineered mutations E78R. A slight asymmetry in this area has Glu89^D^ point toward the core of the bundle (where Glu89^B^ curls up towards the rim) and engage in an H-bond with Q86^C^. Glu 89 was shown to form hydrogen bonds with IKKβ residues S733 and F734^15^ and was reported to interact with the small molecule inhibitor Withaferin in docking studies results^39^.

**Figure 5.**
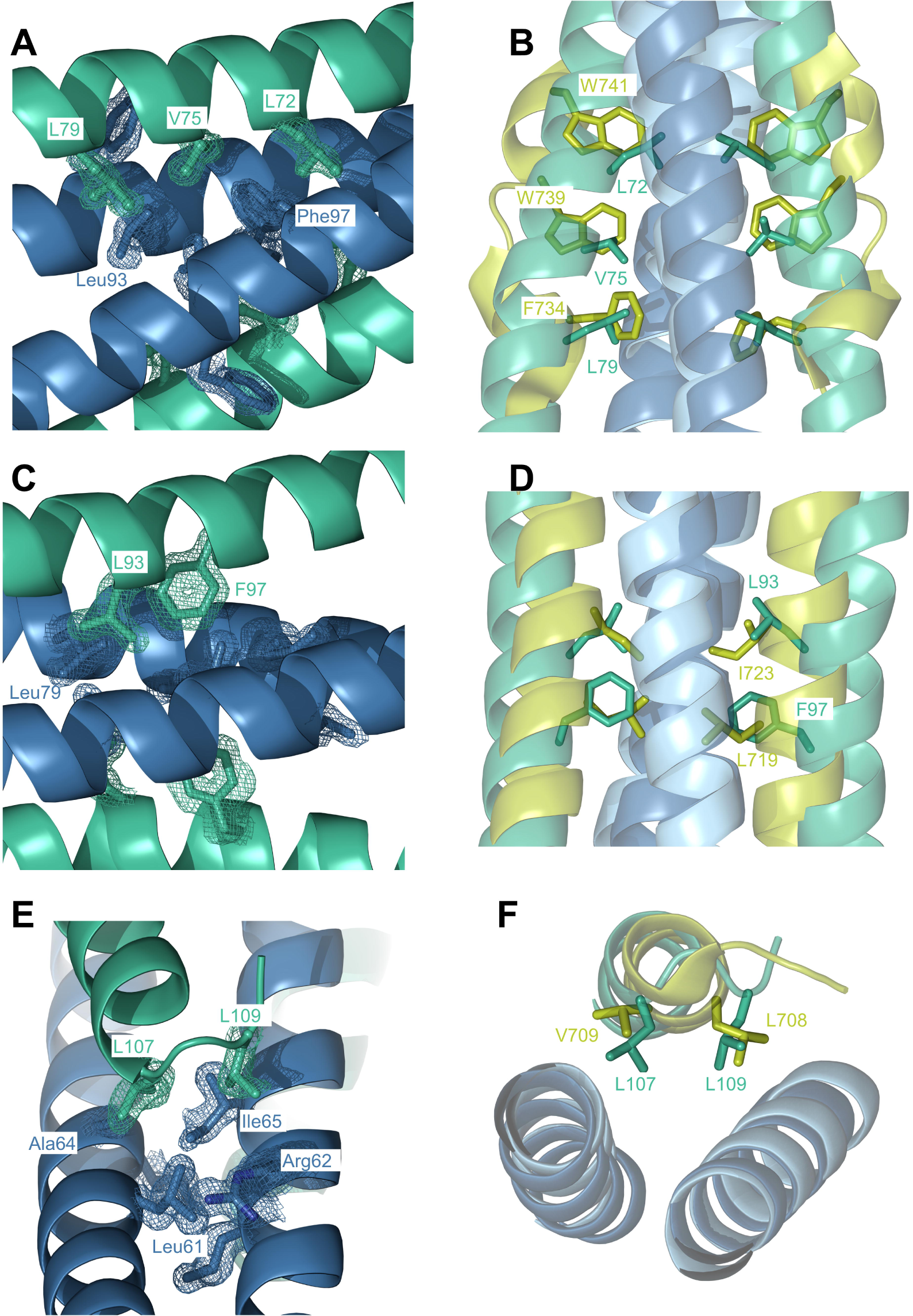
The hot spots for binding are conserved in zipNEMO. The panels illustrate the three hot spots for ligand binding in NEMO: (A, B) hot spot 3; (C, D) hot spot 2; (E, F) hot spot 1. The left panels (A,C,E) show cartoon representations of the zipNEMO structure with the receptor dimer in blue and the pseudo-ligand dimer in mint. Electron density is contoured at 1σ. The right panels (B, D, F) show a superposition of the zipNEMO structure with the NEMO/IKKβ structure (3BRV): NEMO is in light cyan and IKKβ is in pear. Residues of interest are labeled and color coded, using a one-letter code for the “ligands” (greens) and a three-letter code for the receptors (blues).

In the NEMO/IKKβ complex hot spot 2 is defined by IKKβ residues L719, I723, which bind in a pocket formed by NEMO residues Leu72, Arg75, Leu79 (PLIP^40^ analysis of 3BRV). In the zipNEMO dimer the same receptor pocket, residues Leu72, Val75, Leu79 (chains B/D), accommodates pseudo-ligand residues L93 (in the position corresponding to IKKβ L719) and F97 (corresponding to IKKβ I723; Figure 5 C,D). This interaction site creates a binding pocket that is symmetric to the interaction observed for hot spot 3, between chains A/C residues L72, V75, L79 and chains B/D residues Phe92-Phe97. This is coincidental, due to the spacing between the two hot spots in IKKβ (residue clusters 719/723 and 734-741) in the complex structure. The zipNEMO dimer seems to take advantage of the same productive interaction (clusters 72-79 with cluster 92-97) twice, fulfilling both binding hot spots 2 and 3 in NEMO. Again, the R75V mutation seems instrumental in creating a better binding interaction in hot spot 2, by forming a more hydrophobic binding pocket. The hydrophobic and electrostatic interactions in hot spot 2 are similar to hot spot 3, with the exception of Glu74^D^ of hot spot 2, which is involved in coordinating one of the Lanthanide ions.

Finally, hot spot 1 in the NEMO/IKKβ interaction was reported to involve IKKβ residues L708/V709 and to be necessary for stabilizing binding at the N-terminus of the interacting proteins^14^. The pocket for IKKβ residues L708/V709 is formed by NEMO residues 61-64. In the zipNEMO tetramer complex structure residues L107/L109 adopt the same position as IKKβ residues V709/L708 respectively (Figure 5 E,F). The binding pocket is formed by the same NEMO residues Leu61-Ile65. This interaction is observed fully for chain C, which is resolved till residue 110.

We analyzed the high resolution structure of zipNEMO by computational Alanine scanning (Bude Ala Scan^41^, BAlaS), which confirmed the three clusters identified as hot spots for the interaction, involving I71, L72, V75, L79 for hot spot 3, F92, L93, F97 for hot spot 2 and L107, L109 for hot spot 1. These residues have calculated ΔΔG values for Ala substitutions of 1-3 kcal/mol, which is comparable to the experimentally determined ΔΔG values for the IKKβ hot spots of 1-4 kcal/mol^14^. BAlaS also indicates three charged residues as playing an important role in binding (ΔΔG >1): E74 in hot spot 3 (H-bond with Lys96; and symmetrically K96/Glu74 in hotspot 2), R78 in hot spot 3 (H-bond with Glu89, and symmetrically E89/Arg78 in hot spot 2). As a validation, the BAlaS analysis of the NEMO/IKKβ complex structure identifies the same IKKβ hot spot residues as the experimental Ala scanning mutagenesis with comparable ΔΔG values of 1-4 kcal/mol.

### Zipped-NEMO derived peptides bind WT-NEMO

The analysis of the novel head-to-tail, dimer-to-dimer binding of zipNEMO suggests that a non-conserved sequence, unrelated to the sequence of the native ligand IKKβ, could achieve specific binding to NEMO. To probe this hypothesis, we designed two short peptides, based on the zipNEMO sequence and on the analysis of the hot spots for binding and tested them for their ability to bind directly WT-NEMO (Table S1).

The longer peptide, 15 residues, encompasses zipNEMO sequence N69-Q83, and preserves the hydrophobic hot spot 3 residues (L71,L72,V75,L79) as well as residues responsible for most of the H-bond interactions in the complex structure (N69,R73,E74,R78,Q83; Figure S4). A shorter version, 8 residues, L72-L79, preserves the core hydrophobic residues. The 15-mer is predicted to be completely helical by the PEP-FOLD^42^ server, which identifies the lowest-energy conformations. The peptides, labeled with 5-FAM at the N-terminus, bind to our control construct of GST-NEMO(1-196) in the FA assay, with a K_D_ = 44.6 µM (R^2^=0.986, 95% CI: 22.9-66.3) for the 15-mer peptide (Figure S5). The 8-mer peptide shows direct binding but the binding curve was not complete at the highest GST-NEMO(1-196) concentration tested (283 µM). GST-NEMO(1-196) aggregates and precipitates at higher concentration. The binding affinity of the 15-mer peptide is similar to what reported for the NEMO Binding Domain (NBD) peptide (K_D_ = 35 µM by ITC, 18.8-300 µM by SPR^17^, and an IC_50_ = 60 µM by FA in our hands).

## Discussion

In this work we set out to rationally design a NEMO construct that would facilitate structure-based inhibitor development, by enabling structure determination by X-ray crystallography or NMR spectroscopy, and NMR-based inhibitor screening and validation. Modulating protein-protein interactions with small molecule inhibitors entails heightened challenges, due to the usually large and flat interface, the lack of endogenous small molecule ligands and the high affinity between proteins^22,23^. It is often difficult to distinguishing real from artefactual binding, as the drug discovery effort mostly starts from weak hits, and therefore a high degree of validation is recommended for PPIs inhibitors. Structural and biophysical methods can clarify the binding mechanism, identify stoichiometry, highlight artefacts like non-specific inhibition and protein aggregators, identify covalent and allosteric inhibitors and determine which of the two protein partners binds to the inhibitor and the site of binding^22^. Structure-based drug discovery, fragment-based drug-discovery and NMR-based screening were instrumental in the discovery and optimization of the most recent successes in PPI inhibitors development^43,44^, the marketed Bcl-2/Bax PPI modulator Venetoclax^45^ and inhibitors of the XIAP/Caspase9 PPI^46^.

Most of these experimental techniques are not applicable to the full-length NEMO protein, or to the isolated domain of interest, the IKKβ-binding domain. Analysis of NEMO’s sequence (PairCoils2^47^) indicates a decreased propensity to fold as a coiled coil in the region 66-103, the IKKβ binding site, caused by a stutter, (four-residue insert, *abcd* at 72-78)^16^. Irregularities in the fold of coiled-coil proteins often mark interdomain communication or protein-protein interaction sites^48^. The sequence in this region makes the IKKβ binding domain of NEMO an irregular and dynamic coiled coil^16^, causing conformational heterogeneity and instability in the protein and we wondered if we could stabilize a productive binding conformation through mutagenesis while preserving WT-like binding properties. The specific challenge in this effort was to retain a protein small enough to produce high quality NMR spectra, an effort complicated by the long, rod-like structure of coiled coils. The *de novo* design or stabilization of coiled coils has been proposed and tested through a variety of approaches, including sequence optimization, by incorporating favorable hydrophobic residues at *a*/*d* positions^49,50^ and optimizing inter- and intra-strand ionic interactions^51^ (helix and dimer stabilization), helix stabilization through stapling and dimer stabilization through interhelical disulfide bonds or covalent bonds^52–55^.

Recently, a synthetic coiled-coil mimic of the vFLIP binding domain of NEMO was generated by optimizing knob-into-holes packing and using Crosslinked Helix Dimers, where an interhelical salt bridge is replaced by a covalent bond^52,21^. In our lab, the IKKβ binding domain of NEMO was stabilized by ideal coiled-coil adapters fused at the N- and C-termini, yielding a protein that bound IKKβ with a K_D_ comparable with WT-full-length NEMO and yielded an X-ray structure, although the adapters size prevented NMR studies. Using the disulfide stabilization approach, a NEMO construct (1-120 or 44-111) carrying a L107C mutation was reported to stabilize the dimeric coiled-coil structure and bind IKKβ with high affinity^27^. We chose to avoid introducing additional cysteine residues due to the complications we observed in maintaining appropriate redox conditions to preserve the desired disulfide bonds while preventing disulfide-mediated aggregation. We decided instead on a structure-based and computational approach for sequence optimization^32,56^, targeting both hydrophobic residues *a*/*d*, and salt bridge residues *g*/*e*. The results showed that our designed mutations, in absence of additional covalent- or disulfide-enforced dimerization (beyond the native Cys54), are sufficient to produce a stable and folded dimeric coiled coil, where the corresponding WT NEMO sequence was shown to be largely monomeric and with a weak α-helical content^15,27^. The stabilization of the folded structure also corresponded to a 20 fold improvement in binding affinity for IKKβ, in agreement with the hypothesis that the IKKβ binding domain of NEMO is ordered in the dimeric full length protein and that the conformation favored by the engineered mutant is a productive binding conformation. The native-like folding is further confirmed by the fact that a longer construct incorporating the same mutations, zipNEMO(50-196), binds IKKβ with affinity comparable to full length WT-NEMO. As zipNEMO(50-196) still binds IKKβ approximately 40-fold better than zipNEMO(50-113), it is likely that the longer construct provides additional stabilization. It is notable that even a very short NEMO sequence can be stabilized so that to achieve full-length like binding affinity, as in the adapter-stabilized NEMO(50-113), K_D_ = 12 nM^28,29^.

The biophysical and structural data indicate an additive, if not synergic, stabilization of the structure as more mutations were introduced. The N-terminal inter-helix hydrogen bond between Glu50 and Arg55 (group 2: T50E, L55R), verified in the structure, was determinant in stabilizing the structure, improving the helical and coiled-coil content (60 -> 74%). The central Group 1 mutations improved packing with R75V buried within the coil interface and E78R engaged in intra-helix stabilizing i, i+3 H-bonds with His81 and inter-dimer H-bonds with Glu89. The effect of Group 5 mutations is more difficult to interpret, as the residues are at the very C-terminus and mostly disordered or with poor density in the structure. The R106E, E110V mutations were introduced to pin the protein C-terminus, similarly to what achieved by the L107C disulfide bond. Unfortunately, the new apo-NEMO structure (6MI3) showed that both side chains may be solvent exposed and form an intra-chain hydrogen bond, stabilizing the helical structure. An explanation for the stabilizing effect of Group 5 mutations is the possible new intra-chain H-bond between Glu106 and Arg102. It is also possible that additional transient electrostatic or hydrophobic interactions are present in solution, explaining the almost 10-fold improved binding affinity for IKKβ and improved quality in the NMR spectra in NEMO-125.

The final improvement produced by the C76S, C95A mutations and truncation of residues 44-49 confirms that disulfide mediated aggregation is problematic for NEMO constructs’ solution behavior and indicates that the additional disordered N-terminal residues are both unnecessary and deleterious for NMR and X-ray crystallography experiments.

The zipNEMO construct is the first IBD construct to produce ^1^H,^15^N heteronuclear correlation NMR spectra of high quality. Most importantly, we show that the NMR spectra easily detect the chemical and magnetic environment changes as well as the conformational changes the protein experiences upon ligand binding, as in the presence of a 1.1 equivalent of IKKβ(701-745). The sample and experimental conditions are suitable for medium throughput NMR-based screening and the spectral quality will allow for inhibitor validation and determination of binding site within the same experiment (upon backbone resonance assignment).

Additionally, we were able to crystallize and solve the structure of the NMR-screening construct, opening the way to combine screening and complex structure determination by X-ray crystallography. The zipNEMO structure reveals that the fundamental hot spots for the interaction of NEMO with IKKβ are conserved in the interaction between zipNEMO dimers. There is no sequence conservation between IKKβ(701-745) and zipNEMO(50-113). Both protein constructs are helical in the X-ray structure but while zipNEMO is helical throughout, IKKβ unwinds in the region between 732 and 742, to accommodate interactions in hot spot 3 (the linker and NBD peptide regions)^15^. It was previously observed^14^ how the two different kinases IKKα and IKKβ achieve a convergent solution for binding to NEMO: the spatial location of the hot spots is conserved although differences in sequence mean the ligand residues engaged at the hot spot are not conserved. ZipNEMO seems to employ a similar convergent solution for binding, where residues’ spacing and character fulfill the binding hot spots requirements, but this is realized through a different sequence and secondary structure. It is remarkable how a new structural and sequence solution manages to realize a residue side chain presentation so similar to the one of the native ligand IKKβ. It is also interesting how the pseudo-ligand zipNEMO dimer succeeds in fulfilling the most important interactions for binding with a head-to-tail orientation. This can in part be explained by the fact that most interactions are side chain-mediated, not involving backbone atoms H-bonds.

The major difference observed in the zipNEMO structure compared to the NEMO/IKKβ structure is the opening of the binding cleft towards the chains’ C-terminus, to accommodate the pseudo-ligand dimer, longer than IKKβ(701-745). The different degrees of opening of the coiled-coil dimer, as experimentally observed in the apo and the ligand-bound structures (3BRV, 6MI3, 6MI4 and here 8U7C) point to the adaptability of the binding pockets, which depends on the ligand presence and characteristics. A certain degree of flexibility in the IBD is reasonably maintained within the full-length NEMO protein, where it is preceded by the disordered N-terminal sequence (1-50) and followed by the Intervening Domain Region (IVD, residues 112-195). While this region is predicted to fold as a coiled coil (PairCoils2^47^), CD solution studies report that the isolated IVD is a flexible monomer with a low α-helical propensity. Our results reveal that NEMO/IKKβ inhibition can be achieved by a peptide with a new sequence and secondary structure combination, and that the NEMO structure adapts to accommodate the new residues in the same hot spots for binding. As our earlier NEMO structures suggested^16^, the flexibility in the IKKβ binding site should be taken into account when designing peptidic and small molecule inhibitors for the NEMO/IKKβ interaction and reinforces the concept that direct complex structure determination will play a fundamental role in drug design and optimization of NEMO inhibitors. The suggestion that adaptive binding may create an optimal binding pocket for a small molecule inhibitor targeting NEMO should also encourage us to reevaluate the druggability of the pocket on NEMO and the suitability of NEMO as a therapeutic target.

## Acknowledgments

This work was funded by NIH grants R03AR066130, R01GM133844 and R35GM128663. This research used the AMX and FMX beamlines of the National Synchrotron Light Source II, a U.S. Department of Energy (DOE) Office of Science User Facility operated for the DOE Office of Science by Brookhaven National Laboratory under Contract No. DE-SC0012704. We thank the staff and scientists at NSLS II for their outstanding support. This work was supported by bioMT through NIH NIGMS grant P20GM113132. We thank the staff at bioMT for their amazing support. This work was supported by the The Dartmouth Innovations Accelerator for Cancer and by The Elmer R. Pfefferkorn & Allan U. Munck Education and Research Fund Novel and Interactive Grant Initiative; we would like to thank the staff and investigators at the Magnuson Center for Enterpreneurship at Dartmouth and the Dartmouth Cancer Center for their support. AEK was the recipient of a Dartmouth Innovation PhD Fellowship Program.

## Author Contributions

MP, DFM: resources, project administration. MP, DFM, GG, AHB: conceptualization. MP, DFM, MJR: supervision, funding acquisition. MP, AHB, AEK, GG, KAT, MBH, CRA, CCR, JPP:, investigation, formal analysis. MP, AHB, AEK: methodology, visualization, validation. MP: writing – original draft. MP, DFM, MJR: writing – review and editing.

## Declaration of Interest

We, the authors (Pellegrini, Grigoryan, Barczewski), have a patent application related to this work “ENGINEERED NEMO AND USES THEREOF”. U.S. Patent Application No. 63/196,217.

## Methods

### Method Details

#### Construction of E. Coli expression vectors

NEMO vectors incorporating the wild type sequence were subcloned from the Origene plasmid (Cat# RC201743) using standard restriction sites or Agilent’s QuikChange II XL or QuikChange Lightning site-directed mutagenesis kits through a PCR insertion method which precludes restriction enzymes and ligases^57^. Briefly, the DNA primers (GenScript, Piscataway, NJ) were designed to include a distal segment overlapping with the target plasmids and a proximal segment overlapping with the protein construct. In the first step, the proximal segments were used as primers to PCR-amplify DNA segments of target clones with overhangs corresponding to the plasmids. In the second step, the purified PCR products were used for site-directed mutagenesis-based insertion, where the inserts’ distal/overhanging segments defined the insertion points on the target plasmids. The NEMO(44-113) construct was cloned into a pET-16b vector with a TEV cleavage site inserted before the NEMO sequence as described in ^28^. A tryptophan was inserted at the N-terminus of the NEMO construct for UV detection. The final construct has the residues GSW appended before the NEMO sequence as a result of the TEV cleavage and W insertion and is referred to as WT-NEMO(44-113)^28^. This vector was utilized for all subsequent vector’s construction, utilizing site directed mutagenesis and the QuikChange XL II site-mutagenesis kit. Group 1 (R75V, E78R), Group 2 (T50E, L55R), Group 3 (C76D, I71K, L80K), Group 4 (S68N) and Group 5 (R105E, E110V) mutations were introduced in successive rounds. Mutations C76S, C95A, were introduced on the mutant incorporating Group 1, 2, 5 mutations to generate NEMO-125CSCA. Finally, the 6 N-terminal residues of NEMO-125CSCA (EQGAPE corresponding to NEMO residues 44-49) were deleted by site-directed mutagenesis resulting in the final construct called zipNEMO. GST-tagged constructs were cloned into a pGEX-6P-1 vector with an N-terminal GST tag as reported in ^16,28^, and the cleavage site was mutated to a TEV protease cleavage site by side directed mutagenesis (QuikChange mutagenesis kit). Plasmids for GST-zipNEMO(50-196), incorporating the same mutations as zipNEMO, and with a codon-optimized nucleotide sequence were purchased from GenScript. Additional C131A and C167A mutations were introduced into GST-zipNEMO(50-419), using a QuikChange Multi Site-Directed Mutagenesis Kit (Agilent). The expression vector for IKKβ_KKRR_(701–745) was subcloned from Addgene Plasmid #11103 into an in-house engineered pJCC04a vector with a thioredoxin fusion protein and a TEV cleavage site, as described in^28,58^. The K703R, K704R double-lysine mutant (IKKβ_KK/RR_) was generated using site-directed mutagenesis (QuikChange Lightening site-directed mutagenesis kit)^59^. The sequence of all vectors used was confirmed by Sanger sequencing (GeneWiz). New mutagenesis primers used in this study are listed in Table S2.

#### Expression and purification of NEMO constructs

NEMO His-tagged constructs were expressed in BL21 Star (DE3) *E. Coli* cells or Rosetta 2 cells (Novagen). Cells were grown at 37°C to OD_600_ = 0.8 – 1.0 in Terrific Broth (TB) or ZYP-5052 media^60^ before induction with 0.5 mM Isopropyl ß-D-1-thiogalactopyranoside (IPTG), and harvested after 4 hour at 37°C or 20 hours at 20°C. For all proteins, cells were harvested by centrifugation at 4,000-5,000 rpm for 15-20 minutes at 4°C. Cell pellets were resuspended in 25 mL of lysis buffer [50 mM Tris pH 8.0, 250 mM NaCl, 10 mM imidazole, 0.2 mM TCEP, 1 mM phenylmethylsulfonyl fluoride (PMSF), 1 μL DNAse (Benzonase)]. Cells were lysed with a French Press, and the lysate was denatured by adding urea to a concentration of 8 M. Proteins were purified by affinity chromatography on GE Healthcare AktaXpress FPLC systems. The clarified urea lysate was applied to a HisTrap 5 mL HP column (GE Healthcare) and washed using 50 mM Tris pH 8.0, 250 mM NaCl, 8 M urea, 10mM imidazole, 0.2 mM TCEP. Urea was then removed using a refolding buffer consisting of 50 mM Tris pH 8.0, 250 mM NaCl, 10mM imidazole, 0.2 mM TCEP, and the target protein eluted with a 0-500 mM imidazole gradient. Bradford reagent was used to determine the protein concentration. His-tagged TEV protease was added at a ratio of 1 mg of TEV to 10 mg of His-tagged protein for TEV cleavage in dialysis buffer [50 mM Tris pH 8.0, 250 mM NaCl, 0.2 mM TCEP, 5% glycerol] overnight at 4 °C. The cleavage mixture was applied to the HisTrap column again to remove the His tag and TEV, followed by SEC with a Superdex75 16/600 column in different buffers, depending on the protein destined use (see below). Proteins were concentrated using a 50mL Amicon stirred cell concentrator with a 3-5 kDa MWCO disk or with Amicon centrifugal concentrators of the same MWCO. ^15^N labeled proteins were prepared similarly, with the scale up culture grown to an OD_600_ = 3-4, before transferring to M9 media containing ^15^N-NH_4_Cl. Cells were grown for 45 minutes before induction with 0.5 mM IPTG. Cells were harvested after 16 hours of growth at 20°C. ^2^H,^15^N labeled proteins were prepared with a protocol similar to ^61^, adapting cells to increasing concentration of D_2_O in TB media and then switching to D_2_O M9 media containing ^15^N-NH_4_Cl. After induction cells were grown at 37°C for 16 hours before harvesting.

#### Expression and purification of GST-NEMO(1-196)

GST-NEMO(1-196) and GST-zipNEMO(50-196) were produced and purified as described in ^28^. Briefly, cells were grown at 37°C to an OD_600_ = 0.8 – 1.0 in TB or ZYP-5052, induced with 0.5mM IPTG and grown at 37°C for 4 hours, shaking at 220 rpm. Cell pellets were resuspended in 25 ml of 50 mM Tris pH 7.6, 150 mM NaCl, 2 mM TCEP, 10% glycerol, 2 mM MgCl2, 1 μL Benzonase, 0.02% NP-40, 200 μM PefaBloc, and lysed using a French Press. Clarified cell lysate was purified on a GSTrap 4B column (GE Healthcare, washing: 50 mM Tris pH 8, 150 mM NaCl, 5 mM DTT, 1 mM EDTA; elution: 50 mM Tris pH 8, 150 mM NaCl, 5 mM DTT, 1 mM EDTA, 10 mM reduced glutathione), followed by SEC on a Superdex75 16/600 column (GE Healthcare, 50 mM sodium phosphate pH 6.5, 150 mM NaCl, 5 mM DTT. Proteins were concentrated using a stirred cell N_2_ flow Amicon 50 concentrator with a 10 kDa MWCO disk.

#### Expression and purification of zipNEMO with SeMet labeling

ZipNEMO was expressed in B834 (DE3) *E.Coli* cells, methionine auxotrophs. A 250 mL culture was grown at 37°C to an OD_600_ = 3-4. Cells were centrifuged at 1,500 rpm for 10 minutes at 4°C. Cells were then resuspended in M9 minimal media containing 3 g/L of NH_4_Cl and 10 g/L dextrose. Cells were grown for 30 minutes before adding 100 µg/mL of selenomethionine, lysine, phenylalanine, and threonine as well as 50 µg/mL of leucine, isoleucine, and valine. Cells were grown for an additional 30 minutes before induction with 0.5 mM IPTG. Expression and purification followed the same protocol as for the unlabeled constructs.

#### Expression, purification and labeling of IKKβ_KKRR_(701–745)

IKKβ_KKRR_ (701–745) was expressed and purified in the same fashion as NEMO, with denaturing and on-column refolding followed by TEV cleavage and SEC, as described in ^28^. The protein was labeled with fluorescein isothiocyanate (FITC) as described^28^. Purified IKKβ_KKRR_ construct in final purification buffer (20 mM Na phosphate pH 6.5, 50 mM NaCl, 0.2 mM TCEP) was added with 100 mM sodium bicarbonate pH 10.0, and a 100-fold excess of Fluorescein Isothiocyanate (FITC). FITC-labeling was achieved according to the manufacturer’s protocol (CITE) and assessed spectrophotometrically at 280 and 495 nm, using calculations as provided through Thermo Fisher’s FluoReporter FITC Protein Labeling kit.

#### Fluorescence Anisotropy

Direct binding assays were performed using N-terminally FITC-labeled IKKβ_KK/RR_(residues 701–745), using recombinant protein as above or synthetic peptide (GenScript). Measurements were obtained using a Tecan Infinity F500 plate reader at room temperature, and each experiment was run in triplicate. Each well contained 2 mM Tris, 100 mM NaCl, 2 mM DTT, pH 8.0, 0.1 mg/mL IgG and 0.5 mM Thesit, 30 nM FITC-labeled tracer peptide and NEMO proteins at varying concentrations. Plates were shaken for 20 seconds and incubated for 15 minutes at room temperature, prior to data collection. The anisotropy values, FA, were fitted by nonlinear least-squares regression to the modified quadratic binding equation shown below using MATLAB (R2023a, The MathWorks, Inc., Natick, Massachusetts, United States):

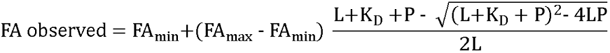

where FA_min_ is the anisotropy value of the reporter in absence of NEMO, FA_max_ is the maximum anisotropy value at saturating NEMO concentrations, L and P are the total concentrations of reporter and NEMO, and K_D_ is the dissociation constant. The fitting is carried out on the raw FA values, while the figures are displayed on a normalized FA scale of 0 to 1 for ease of comparison of the different constructs.

#### Cross-linking experiments

Cross-linking experiments utilized samples of zipNEMO in 50 mM sodium phosphate, pH 6.5, 250 mM NaCl, 0.2 mM TCEP, and in 25 mM HEPES, pH 6.5, 424 mM NaCl, 0.2 mM TCEP, at a concentration of 100-280 µM. The reference sample WT-NEMO(44-113) was 100 µM in 20mM sodium phosphate pH 6.5, 50mM NaCl, 0.2 mM TCEP. Cross-linking was carried out using bis(sulfosuccinimidyl)glutarate (BS2G, ProteoChem; 7.7 Å) and bis(sulfosuccinimidyl)suberate (BS3, ThermoFisher; 11.4 Å) homobifunctional protein crosslinkers with spacer arms of the indicated length. Frozen stocks of protein in 50 mM Na phosphate pH 6.5, 250 mM NaCl, 0.2 mM TCEP were thawed were dialyzed into 25 mM HEPES pH 6.5, 424 mM NaCl, 0.2 mM TECEP in a micro dialysis cassette at room temperature. The protein was then concentrated in the respective buffers to 100-280 µM. 50 µL aliquots were prepared for each reaction. Just before the reaction, 2 mgs each of BS3 or BS2G were dissolved in the Na phosphate buffer and the HEPES buffer, respectively, for a final concentration of 50 mM. Crosslinkers were added to the aliquots to a final concentration of 5 mM. The reaction was allowed to proceed for 30 minutes at room temperature, then the reaction was quenched with 50 mM tris and allowed to incubate at room temperature for 15 minutes. The samples were then prepared for gel electrophoresis with the addition of SDS-PAGE Protein Loading Buffer (5x) with and without 10 mM DTT present to assess the presence of disulfide-mediated oligomers in each sample. Controls of unreacted protein were also prepared with and without DTT. The samples were boiled for 10 minutes, quickly vortexed, then spun down at 10,000 RPM for 5 minutes. 10 µL of each sample was loaded onto a 12% sucrose polyacrylamide gel and run at 145 V for 55 minutes using SeeBlue Plus as molecular weight standards (Invitrogen).

#### SEC-MALS

SEC was performed using a Superdex75 10/300 column (GE Healthcare) in 20 mM sodium phosphate (pH 6.8) and 150 mM NaCl, with an AKTAPure instrument (Cytiva) coupled to an Optilab refractive index detector (Wyatt) and light-scattering detector (Wyatt MiniDawn Tristar). The molar mass was determined using the ASTRA Software. The system was calibrated using BSA assuming isotropic light scattering. The protein samples were 1.5 mg/ml.

#### NMR spectroscopy

All NMR experiments were carried out on either a Bruker Avance 700 MHz or 600 MHz spectrometer equipped, respectively, with 5 mm and 1.7 mm TCI cryoprobes. Spectra were acquired at 308 K using 50-300 µM samples of NEMO (unlabeled, ^15^N or ^2^H^15^N labeled) in NMR Buffer consisting of 50 mM Na phosphate pH 6.5, 250 mM NaCl, 0.2 mM TCEP, or 25 mM HEPES pH 6.5, 50 mM NaCl, 0.2 mM TCEP. IKKβ(701-745) binding was assessed on a sample containing 100 µM zipNEMO and a 1.1 fold excess of peptide in PBS. The control experiment contained 100 µM zipNEMO in PBS. 1D ^1^H experiments utilized water suppression by excitation sculpting, 2D ^1^H, ^15^N-HSQC^62,63^ and 2D ^1^H, ^15^N-TROSY spectra were acquired with standard Bruker’s pulse sequence. Each 2D spectrum was recorded as a complex data matrix with 1024 x 128 or 256 points and 8-64 scans per FID. Data were processed using Topspin 3.5 or 4.0.8.

#### Circular Dichroism

Coiled-coil character and construct stability were assessed by circular dichroism (CD). All intermediate constructs were analyzed at concentrations of 5, 10, 15, 20 and 25 µM in 20 mM sodium phosphate, pH 6.5, 50 mM NaCl, 0.2 mM TCEP, unless otherwise noted. The final construct, zipNEMO was examined at 15, 19 and 23 µM in 50 mM sodium phosphate, pH 6.5, 250 mM NaCl, 0.2 mM TCEP and 20 mM HEPES pH 6.5, 424 mM NaCl, 0.2 mM TCEP. CD data was acquired in a 0.1 cm cell, over 3-10 accumulations from 198-205 nm to 250-255 nm at 20°C at 100 nm/min, and with a 1 nm bandwidth. Melting curves were obtained through variable temperature scans at fixed wavelength (208 and/or 222 nm) from 20°C to 90-95°C at a 1°C/minute ramp rate. The melting temperature T_m_ for each protein was estimated from the maximum of a plot of the first derivative of θ_222_ against temperature. Helical content was estimated from the CD spectra using K2D3^64^.

#### Protein Crystallization of NEMO-EEAA and NEMO-I65M

Screening utilized MRC2 96 well sitting drop plates and a Formulatrix NT8 drop dispenser, with 200 nL drops at a 1:1 and 1:2 protein solution to mother liquor ratio. Fine screens were produced using a Formulatrix Formulator. Initial crystals were obtained with a Morpheous II screen (Molecular Dimensions) using a protein concentration of 140 µg/ml in 20 mM HEPES, 424 mM NaCl, 2 mM DTT, pH 6.5. An additive screen with Morpheus® II - 0.02M Lanthanoides Mix was utilized to improve crystallization. Final crystals were obtained with seeding in 40% Morpheus precipitant mix 7, 0.1 M Morpheus buffer system 4 pH 6.5, 0.00266 M Lanthanoides Mix, 1 M L-Proline (Millipore Sigma). The crystals were cryoprotected by adding 2M L-Proline to the well prior to looping, and flash-frozen in liquid nitrogen. All crystal imaging used a Formulatrix Rock Imager 1000. Crystals of the SeMet zipNEMO were obtained in identical conditions.

#### Data Collection and Structure Determination

SAD diffraction data were collected in cryogenic conditions at the FMX beamline at the National Synchotron Light Source II (NSLS II), Brookhaven National Laboratory at the Se edge (λ=0.979106 Å). Four data sets were acquired from different, non-overlapping spots within the same crystal. The data were processed using XDS^65^ manually or by the program AutoPROC^66^ (Global Phasing Inc.) and analyzed and merged with BLEND^67–69^ within the CCP4 suite^70^. The final blended data was anisotropically truncated using the STARANISO^71^ server (http://staraniso.globalphasing.org/cgi-bin/staraniso.cgi) to define anisotropic diffraction cut-off surfaces without the assumption of an ellipsoid shape, and to apply an anisotropic correction factor to the amplitudes with Bayesian estimation^72^. The anisotropic diffraction cut-off was defined by a locally averaged value of I/σ(I) > 1.2, as suggested by the server. The anisotropic truncation of the data, resulted in new resolution limits of 1.878 Å, 1.395 Å and 1.671 Å along the a*, b* and c* axis. R_free_ flags were defined (6.32% of reflections) using Phenix’s Reflection File Editor. Anomalous scatterers were identified by the HySS (Hybrid Substructure Search) submodule of the Phenix package. The top scoring consensus model contained 8 scatterers with occupancies between 2.97-0.60, log-likelihood gain (LLG) score of 753 and correlation coefficient of 0.384 (at a resolution of 3 Å). The consensus model with 8 scatterers sites was used as input in Phenix Autosol. The top solution of Autosol had 6 refined heavy atom sites, 175 built residues and the following scoring parameters: overall score of 61.8, skew of 0.32, and figure of merit of 0.59, which indicated a correct solution. The model was progressively built by successive cycles of manual building within COOT^73^ and refinement with Phenix.refine. The anomalous scatterer did not correspond in number or coordinates to the SeMet residues in the protein. Analysis of buffer composition suggested that the signal could be due to the Lanthanide ions used as additives in the crystallization buffer. Lanthanides are reported to bind ionically via oxygen-containing sidechains and carbonyls, provide strong anomalous scattering at their L_III_ absorption edge, and have been successfully used for experimental phasing of macromolecules^74,75^. They maintain strong anomalous scattering even at wavelength far from their L_III_ absorption edge^76–78^. Six L_III_ ions were modeled in the spherical electron density. The ions were assigned as three Tb, one Er and three Yt, based on the electron-density and B-factors. Five of them showed clear coordination by 2-[bis-(2-hydroxy-ethyl)-amino]-2-hydroxymethyl-propane-1,3-diol (bis-tris-propane) molecules (BTB), present in the crystallization buffer, as previously observed^79^, which were manually built using COOT. Parameters for bis-tris-propane were obtained from the PDB databank (BTB.cif) and geometry restraint information for refinement was generated by eLBOW, within Phenix. The L_III_ ions and their coordination in the final coordinates were checked using the “CheckMyMetal (CMM): Metal Binding Site Validation Server”^80^. The final coordinates include 139 water molecules, two Prolines, 5 molecules of pentane-1,5-diol and two short PEG fragments form the crystallization conditions. Data collection statistics and refinement parameters are given in Table 2. Pocket analysis in the structures utilized PLIP^40^, interaction analysis utilized Bude Ala Scan^41^, peptide sequence analysis utilized PEP-FOLD^42^. The figures were generated using PyMOL^81^. The color scheme utilized in all figures and graphics is a color blind friendly^82–84^ light qualitative colour scheme (Paul Tol, https://personal.sron.nl/~pault/#sec:qualitative).

### Quantification and Statistical Analysis

First derivatives for the determination of T_m_ from CD thermal melting curves were calculated using Microsoft Excel. Data for the K_D_ determination were fitted using MATLAB, errors are reported as 95% confidence bounds.

## Supplementary Information

Figures S1–S5, Tables S1-S2.

## Notes

### Competing Interest Statement

A my E. Kennedy: Oxford Oxford Cryosystems, Hanover, NH, US
Adam H. Barczewski: Incyte, Wilmington, DE, US
Gevorg Grigoryan Generate Biomedicines, Inc., Arlington, MA, US
We, Pellegrini, Grigoryan, Barczewski, have a patent application related to this work
"ENGINEERED NEMO AND USES THEREOF". U.S. Patent Application No.
63/196,217

