## Supplementary Material for "The structure of a NEMO construct engineered for screening reveals novel determinants of inhibition"

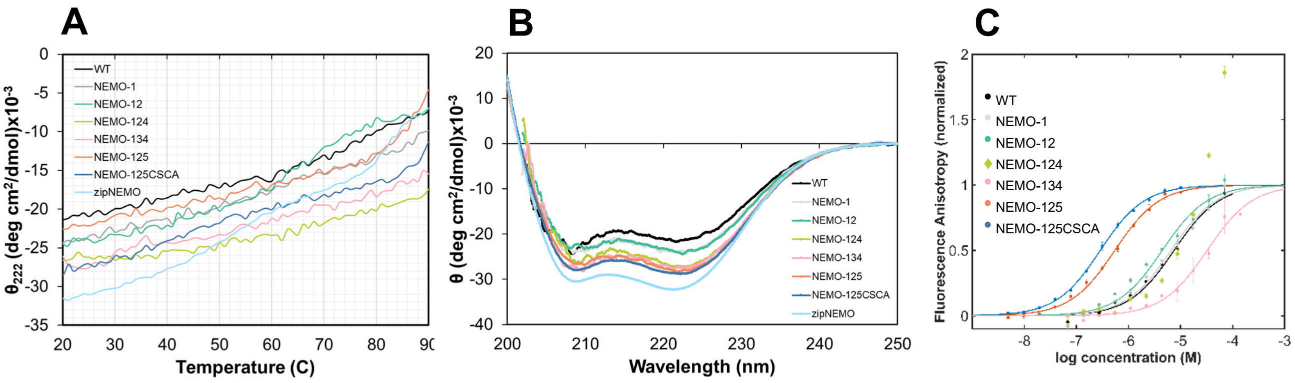


**Figure S1.** Evaluation of the engineered NEMO mutants, related to Figure 2. (A) Thermal denaturation monitored by CD at 222 nm. (B) Overlay of the CD spectra for the engineered NEMO constructs, showing the stabilization of structure and coiled-coil content compared with WT NEMO(44-113). (C) Binding affinity of the engineered NEMO constructs for FITC-IKKβ$_{KKRR}$(701-745) by Fluorescence Anisotropy, compared with WT NEMO(44-113); lines represent the curve fitting. Error bars represent the standard deviation of 3 repeats. NEMO-124 (pear green) displayed aggregation behavior and the curve was not fitted.

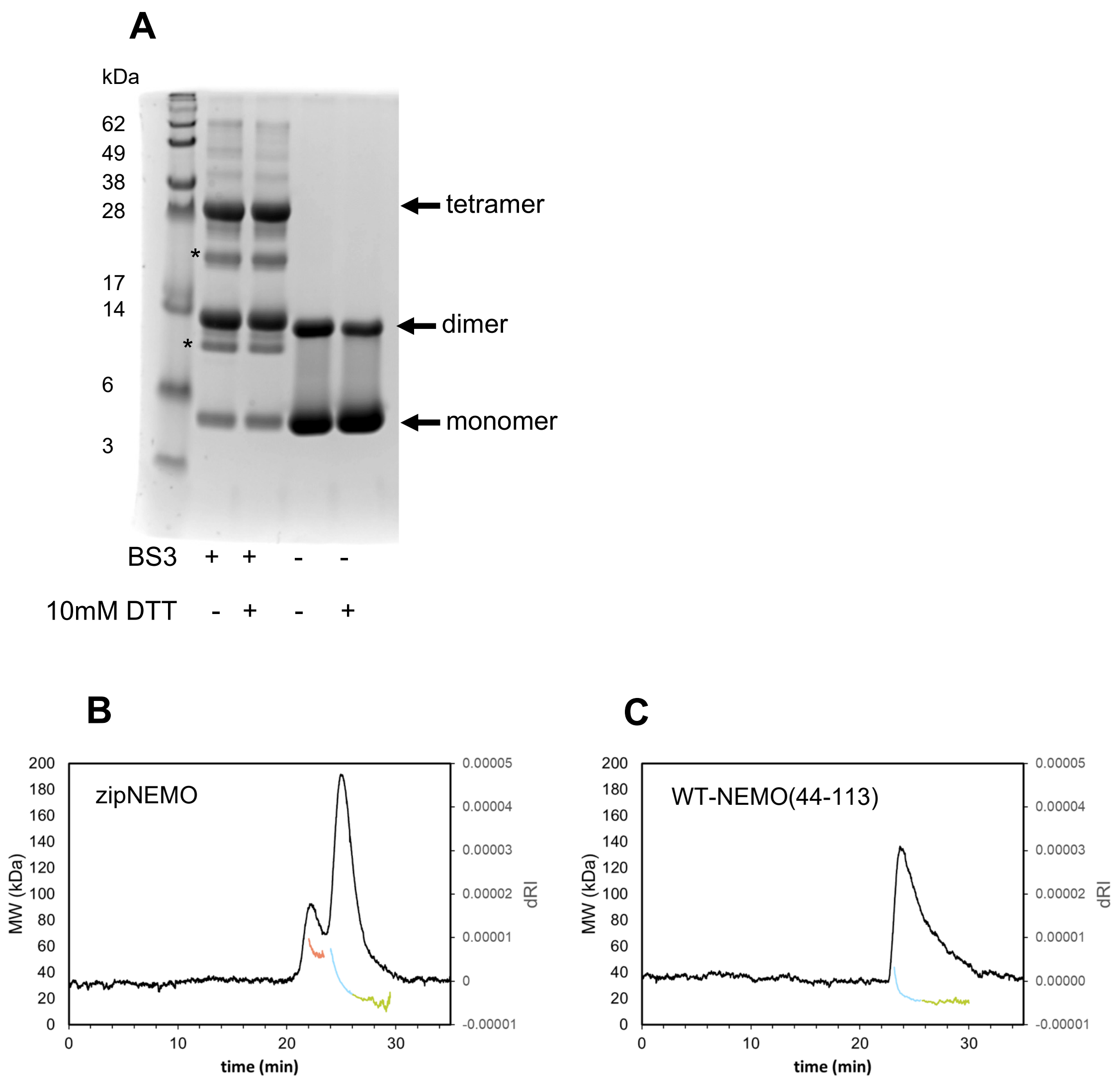


**Figure S2**: Characterization of the oligomerization state of zipNEMO, related to Figure 2. (A) cross-linking experiment using BS3 linker (11.4 Å). zipNEMO is 100 µM in HEPES buffer. DTT presence is indicated. * indicate protein degradation products during the experiment incubation time. (B) SEC-MALS profile of zipNEMO. (C) SEC-MALS profile of WT-NEMO(44-113). The molar mass is plotted at the base of the peak. NOTE: the aggregated higher MW peak in zipNEMO (orange) appears only on thawed frozen samples: only the dimer/tetramer peak appears on the SEC profile of the fresh protein.

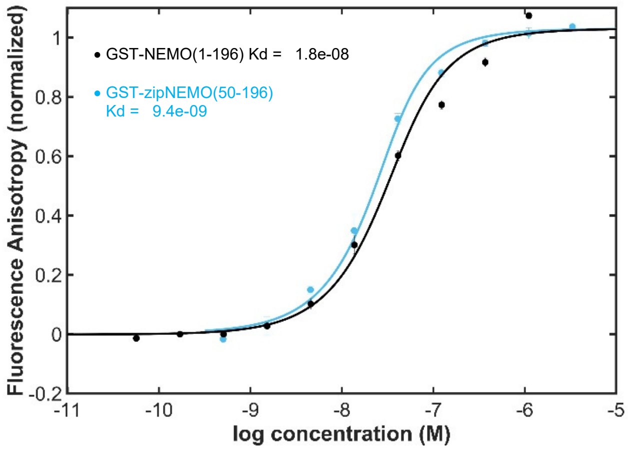


**Figure S3.** Binding affinity of GST-zipNEMO(50-196) for FITC-IKKβ$_{KKRR}$(701-745) by Fluorescence Anisotropy, related to Figure 2.

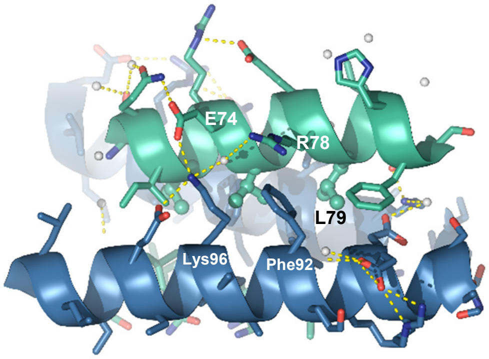


**Figure S4.** zipNEMO-derived 15-mer peptide design, related to figure 5. The figure illustrates the putative interactions of the 15-mer peptide (in mint: $N^{69}$-Q-I-$L^{72}$-R-E-$V^{75}$-S-E-R-$L^{79}$-L-H-F-$Q^{83}$) with NEMO (in blue) as identified from the zipNEMO structure.

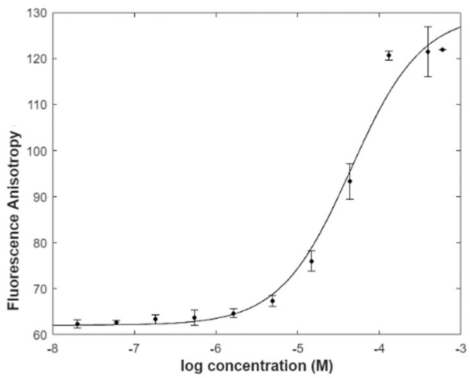


**Figure S5.** Binding affinity of the FAM-labeled 15-mer peptide for GST-NEMO(1-196) by Fluorescence Anisotropy, related to “Zipped-NEMO derived peptides bind WT-NEMO”.

**Table S1** Sequence of the zipNEMO-derived synthetic peptides.

| Peptide | Sequence | N-terminal modification | MW |
|---|---|---|---|
| **8-mer** | L72-R-E-V-S-E-R-L79 | 5-FAM-Ahx | 1472.60 |
| **15-mer** | N69-Q-I-L-R-E-V-S-E-R-L-L-H-F-Q83 | 5-FAM | 2240.44 |

**Table S2** Mutagenesis primers.

| | |
|---|---|
| Group 1 | 5′ - G AGC AAC CAG ATT CTG CGG GAG GTC TGC GAG CGC CTT CTG CAT TTC CAA GCC AGC CAG - 3′<br>3′ - CTG GCT GGC TTG GAA ATG CAG AAG GCG CTC GCA GAC CTC CCG CAG AAT CTG GTT GCT C - 5′ |
| Group 2 | 5′ - GAA CAG GGC GCT CCT GAG GAA CTC CAG CGC TGC CGG GAG GAG AAT CAA GAG CTC CGA G - 3′<br>3′ - C TCG GAG CTC TTG ATT CTC CTC CCG CAA GCG CTG GAG TTC CTC AGG AGC GCC CTG TTC - 5′ |
| Group 4 | 5′ - CAA GAG CTC CGA GAT GCC ATC CGG CAG CTC AAC CAG ATT CTG CGG GAG CGC TGC - 3′<br>3′ - GCA GCG CTC CCG CAG AAT CTG GTT GAG CTG CCG GAT GGC ATC TCG GAG CTC TTG - 5′ |
| Group 3+4 | 5' - GAG CTC CGA GAT GCC ATC CGG CAG CTC AAC CAG AAG CTG CGG GAG GTC GAC GAG CGC CTT AAG CAT TTC CAA GCC AGC CAG AGG GAG GAG - 3'<br>3' - CTC CTC CCT CTG GCT GGC TTG GAA ATG CTT AAG GCG CTC GTC GAC CTC CCG CAG CTT CTG GTT GAG CTG CCG GAT GGC ATC TCG GAG CTC - 5' |
| Group 1+4 | 5′ - CAA GAG CTC CGA GAT GCC ATC CGG CAG CTC AAC CAG ATT CTG CGG GAG GTC TGC GAG CGC CTT CTG CAT TTC CAA GCC AGC CAG GAG G - 3′<br>3′ - C CTC CTG GCT GGC TTG GAA ATG CAG AAG GCG CTC GCA GAC CTC CCG CAG AAT CTG GTT GAG CTG CCG GAT GGC ATC TCG GAG CTC TTG - 5′ |
| Group 5 | 5'- GAG GCC AGG AAA CTG GTG GAG GAA CTC GGC CTG GTG AAG ac GAG TGA GAT CCG GCT-3′<br>3′- AGC CGG ATC TCA CTC GAG TTC CAC CAG GCC GAG TTC CTC CAC CAG TTT CCT GGC CTC-5' |
| C76S | 5' - AGC AAC CAG ATT CTG CGG GAG GTC AGC GAG CGC CTT CTG CAT TTC CAA- 3'<br>3' -TTG GAA ATG CAG AAG GCG CTC GCT GAC CTC CCG CAG AAT CTG GTT GCT- 5' |
| C95A | 5' - CAG AGG GAG GAG AAG GAG TTC CTC ATG GCC AAG TTC CAG GAG GCC AGG AAA CTG-3'<br>3' - CAG TTT CCT GGC CTC CTG GAA CTT GGC CAT GAG GAA CTC CTT CTC CTC CCT CTG - 5' |
